# Asynchronous nuclear cycles in multinucleated *Plasmodium falciparum* enable rapid proliferation

**DOI:** 10.1101/2021.04.15.440016

**Authors:** Severina Klaus, Patrick Binder, Juyeop Kim, Marta Machado, Charlotta Funaya, Violetta Schaaf, Darius Klaschka, Aiste Kudulyte, Marek Cyrklaff, Vibor Laketa, Thomas Höfer, Julien Guizetti, Nils B. Becker, Friedrich Frischknecht, Ulrich S. Schwarz, Markus Ganter

## Abstract

Malaria-causing parasites proliferate within erythrocytes through schizogony, forming multinucleated stages before cellularization. Nuclear multiplication does not follow a strict geometric 2^n^ progression and each proliferative cycle produces a heterogeneous number of progeny. Here, by tracking nuclei and DNA replication, we show that individual nuclei replicate their DNA at different times, despite residing in a shared cytoplasm. Extrapolating from experimental data using mathematical modeling, we demonstrate that a limiting factor must exist that slows down the nuclear multiplication rate. Indeed, our data show that temporally overlapping DNA replication events were significantly slower than partially or non-overlapping events. Our findings suggest an evolutionary pressure that selects for asynchronous DNA replication, balancing available resources with rapid pathogen proliferation.

## Introduction

Malaria is caused by unicellular eukaryotic parasites of the genus *Plasmodium*, with *Plasmodium falciparum* (*P. falciparum*) contributing the most to global malaria-associated morbidity and mortality (1). Disease severity is directly linked to asexual parasite proliferation inside erythrocytes during the blood-stage of infection, which can produce >10^12^ parasites per patient (2). Each asexual proliferative cycle gives rise to approximately 20 daughter cells within 48 hours via a division process called schizogony (3). During early schizogony, nuclei are thought to divide asynchronously (4–6), while schizogony concludes with a relatively synchronous final round of nuclear division that coincides with daughter cell assembly (7, 8). Here, we reveal the reasons driving this unusual multiplication by combining different imaging modalities with computer simulations to understand how DNA replication and nuclear divisions are organized.

## Results

Two distinct models have been proposed to describe the chronology of DNA replication and nuclear division events during *P. falciparum* proliferation in the blood stage (Fig. 1A). Model 1 assumes several rounds of DNA replication prior to nuclear divisions (9, 10), predicting that cells with a single nucleus can have a large distribution of DNA contents. Model 2 proposes alternating rounds of DNA replication and nuclear divisions (6, 11), predicting a gradual increase of both number of nuclei per cell and total DNA content. In addition, this model also assumes that between divisions, nuclei are independent. To test the predictions of both models, we quantified the total DNA content (*C*) (12, 13) and the number of nuclei per parasite (Fig. 1B and Fig. S1A). This showed a positive correlation between DNA content and number of nuclei, supporting model 2. Additionally, the total DNA content never exceeded 2*C* per nucleus, suggesting that the DNA content of individual nuclei alternates between 1*C* and 2*C*.

**Fig. 1.**
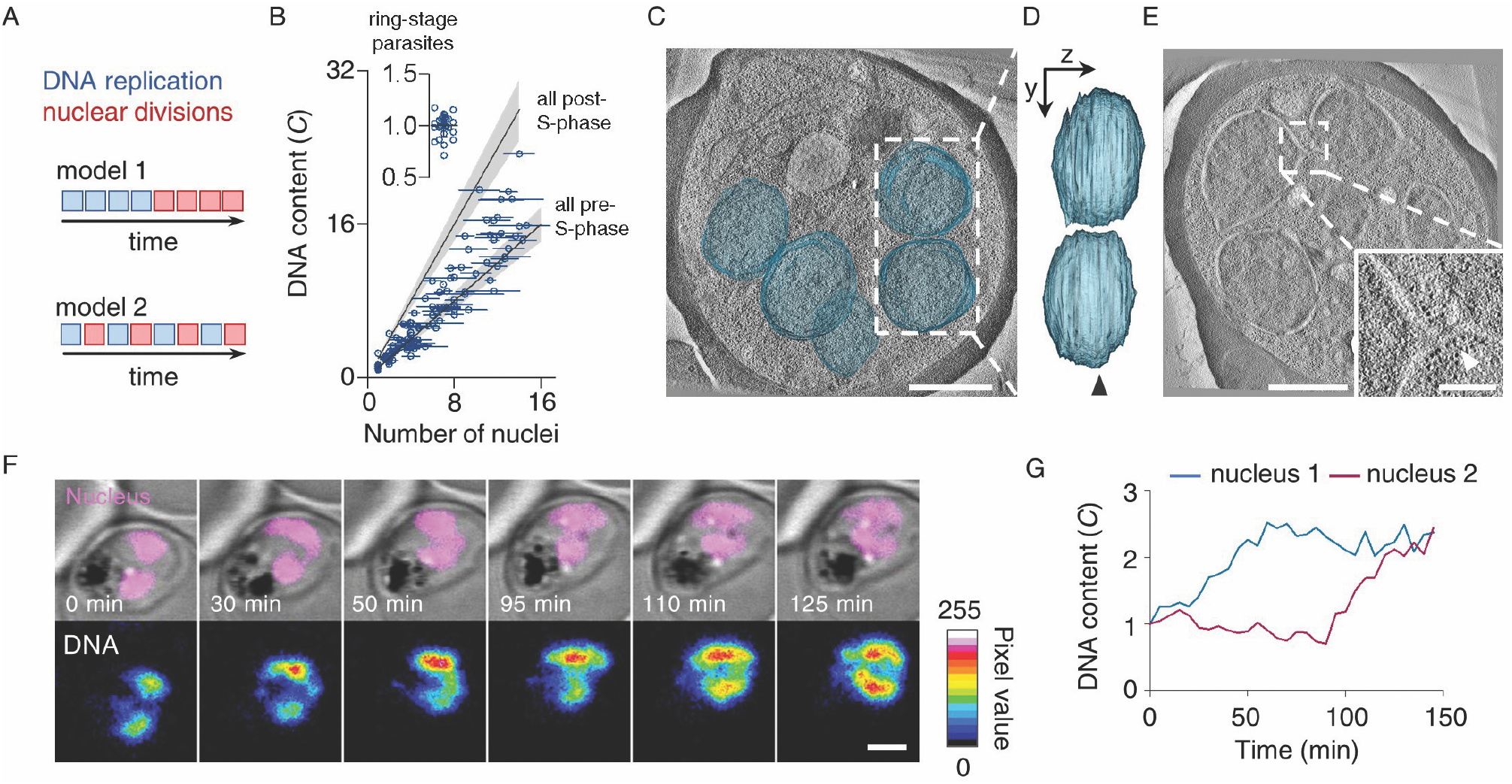
*P. falciparum* proliferates through consecutive rounds of asynchronous DNA replications and nuclear divisions. (A) Schematic of two models proposing the mode of *P. falciparum* proliferation in the blood stage of infection. (B) Gradual increase of the total DNA content and the number of nuclei of *P. falciparum* supports model 2. The DNA content (C) was normalized to pre-S-phase ring-stage parasites (insert), defined as 1. Horizontal bars, standard deviation; gray lines, expected *C*-values for parasites with all nuclei pre- or post-S-phase; gray bands, propagated error (standard deviation) of ring-stage measurements. (C) Electron tomogram, overlayed with 3D-segmented inner nuclear membranes (blue); bar, 1 µm, movie S1. (D) Side view of nuclear volumes showed no connection (90° rotation around the y axis); arrowhead, tomogram plane shown in C. (E) Electron tomogram of connected nuclei; bar, 1 µm; inset highlights the connection (arrowhead); bar, 250 nm. (F) Time-lapse microscopy of a reporter parasite stained with the DNA dye 5-SiR-Hoechst showed asynchronous DNA replication in sister nuclei; bar, 2 µm; movie S3. (G) Quantification of the DNA content of the nuclei shown in F.

To test if nuclei are separate compartments, we recorded three-dimensional electron tomographic views of cell parts containing several entire nuclei (Fig. 1C, D and Fig. S1B, C). Although the nucleoplasm of adjacent nuclei was separated by as little as 75 nm (Fig. S1D, E), most nuclei appeared as independent compartments with clearly discernible nuclear envelopes and ribosomes filling the cytoplasmic gap. In only one out of eight analyzed cells, we recorded a narrow bridge interconnecting two nuclei, which appeared to be completing nuclear division (Fig. 1E). Together these data support a mode of *P. falciparum* proliferation that consists of alternating rounds of DNA replication and nuclear divisions before cellularization. Although nuclear divisions lack synchronization (4–6), it is unclear whether DNA replication in pairs of sister nuclei is synchronized. Employing time-lapse live-cell microscopy, we found that the DNA content increased at different times in sister nuclei, indicating that DNA replications can occur asynchronously, i.e., onset and end of S-phases are desynchronized (Fig. 1F, G and Fig. S2A–D).

To understand how asynchronous DNA replications are orchestrated, we investigated the localization of the DNA replication machinery, using the *P. falciparum* proliferating cell nuclear antigen (PCNA) 1 as a proxy. PCNA is a critical co-factor of DNA polymerases and serves as a hub for many other components of the replication fork (14). Using correlative light and electron microscopy, we found that, in contrast to previous reports (15, 16), ectopically expressed PCNA1::GFP localized unequally in nuclei of the same parasite, with only some nuclei showing distinct PCNA1::GFP foci (Fig. 2A and Fig. S3, S4A, B). Additionally, time-lapse imaging revealed a dynamic localization and transient accumulation of PCNA1 in changing subsets of nuclei (Fig. 2B and Fig. S4C). An increasing nuclear PCNA1::GFP signal was accompanied by a decreasing cytosolic signal and *vice versa* (Fig. 2B, C and Fig. S4D), suggesting that nuclei access a common cytoplasmic pool of PCNA1. Moreover, nuclear accumulation of PCNA1::GFP coincided with a duplication of the DNA content in the same nuclei (Fig. 2D and Fig. S4E, F). This allowed us to track individual DNA replications and nuclear division events over time in a given cell (Fig. 2B, Fig. S5). We defined a nuclear cycle as the total time from the start of an S-phase (S_i_) until the start of the ensuing S-phases in the daughter nuclei (S_i+1,1_ or S_i+1,2_) (Fig. 2E). Additionally, we assigned branches of the nuclear lineages by the onset of S-phase, such that DS_i,1_ ≤ DS_i,2_ etc. The timing of events varied markedly between individual parasites (Fig. S5) and all phases of the first nuclear cycle were longer on average than the corresponding phases of the ensuing cycle (Fig. 2 F–H and Fig. S5). The durations of S-phases and SD-phases were statistically indistinguishable between nuclei of the same generation (Fig. 2F, G) and S-phase duration in the third generation remained similar to those of the second generation (Fig. 2F). Thus, the asynchrony of DNA replications is largely introduced by the times between S-phase events.

**Fig. 2.**
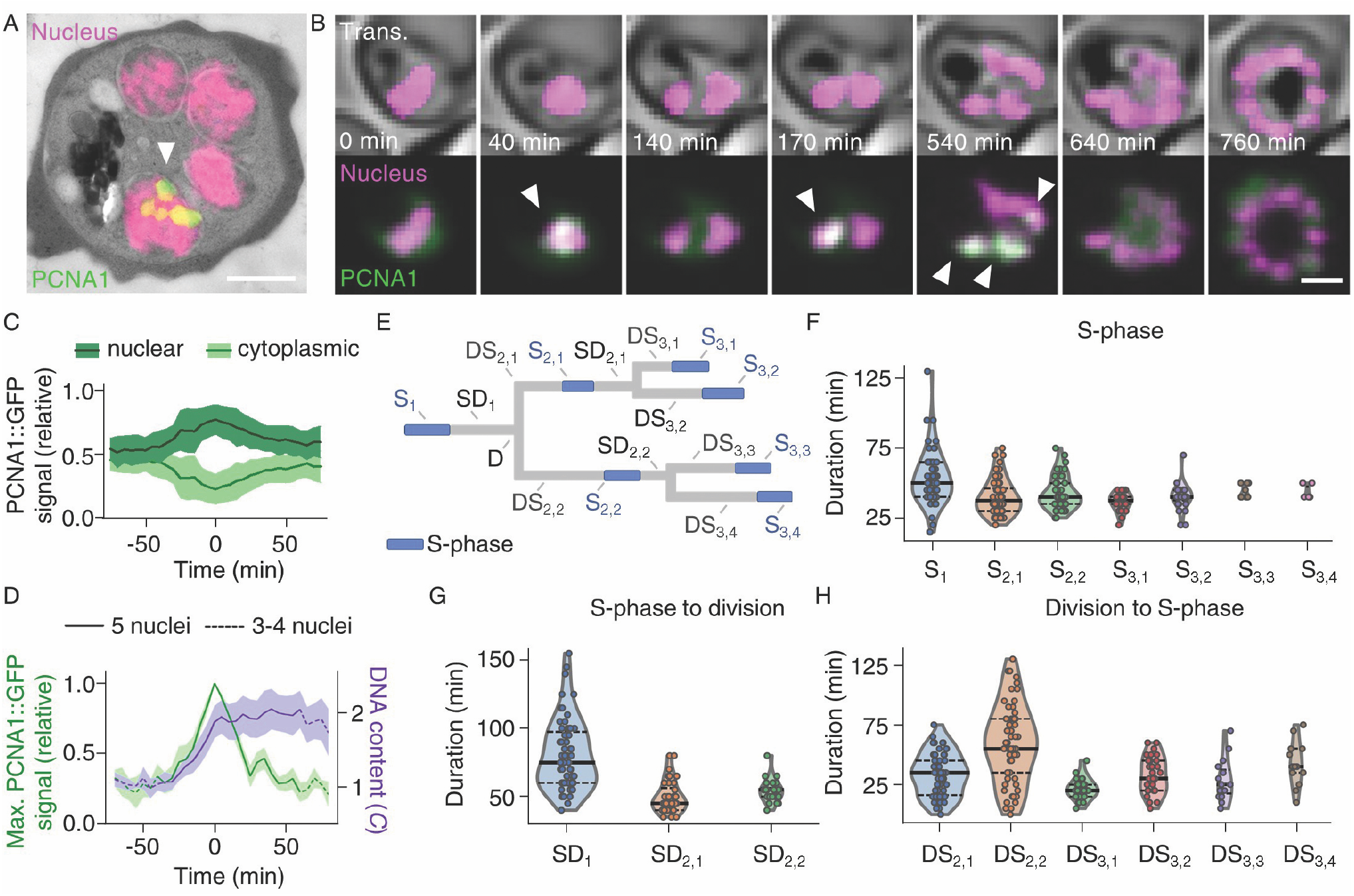
Heterogenous accumulation of PCNA1 among nuclei permits quantification of nuclear multiplication dynamics. (A) Correlative light and electron microscopy showed heterogeneous accumulation of PCNA1 among *P. falciparum* nuclei; bar, 1 µm; arrowhead, PCNA1::GFP focus. (B) Time-lapse microscopy showed dynamic and transient accumulation of PCNA1; bar 2 µm; arrowheads, nuclear PCNA1::GFP accumulation; movies S4, S5. (C) Nuclear accumulation of PCNA1 coincided with a depletion of the cytosolic pool; lines, average (n = 4); bands, standard deviation. (D) Nuclear PCNA1::GFP accumulation caused a peak in signal intensity, coinciding with DNA content duplication; solid lines, average; bands, standard deviation. (E) Schematic illustrating the nuclear cycle phases quantified in F, G, H; S, S-phase; SD, end of S-phase to nuclear division; D, nuclear division; DS, nuclear division to start of S-phase; subscripts (i;j) nuclei within generation i, ordered by their DS phase duration, such that DS_2,1_<DS_2,2_; DS_3,1_<DS_3,2_; and DS_3,3_<DS_3,4_. (F-H) Duration of nuclear cycle phases: S (F), SD (G) and DS (H); solid lines, median; dashed lines, quartiles.

Later rounds of S-phase and nuclear division could not be confidently analyzed by live-cell microscopy due to the spatial proximity of nuclei (Fig. S1D, E). Hence, we aimed to extrapolate the dynamics of nuclear multiplication by mathematically modeling it as a branching process. Our computer simulations were parameterized using the observed distributions of the duration of S-phases and the intervals between S-phases of the first and second nuclear cycles (i.e., S_1_, SD_1_+DS_2_, S_2_, and SD_2_+DS_3_) (Fig. 2F–G and Fig. S6A–C). As the durations of the first and second nuclear cycles (i.e., start S_1_ to start S_2_ versus start S_2_ to start S_3_) showed no significant correlation (Fig. 3A), we did not include the inheritance of factors that facilitates nuclear multiplication in the model.

**Fig. 3.**
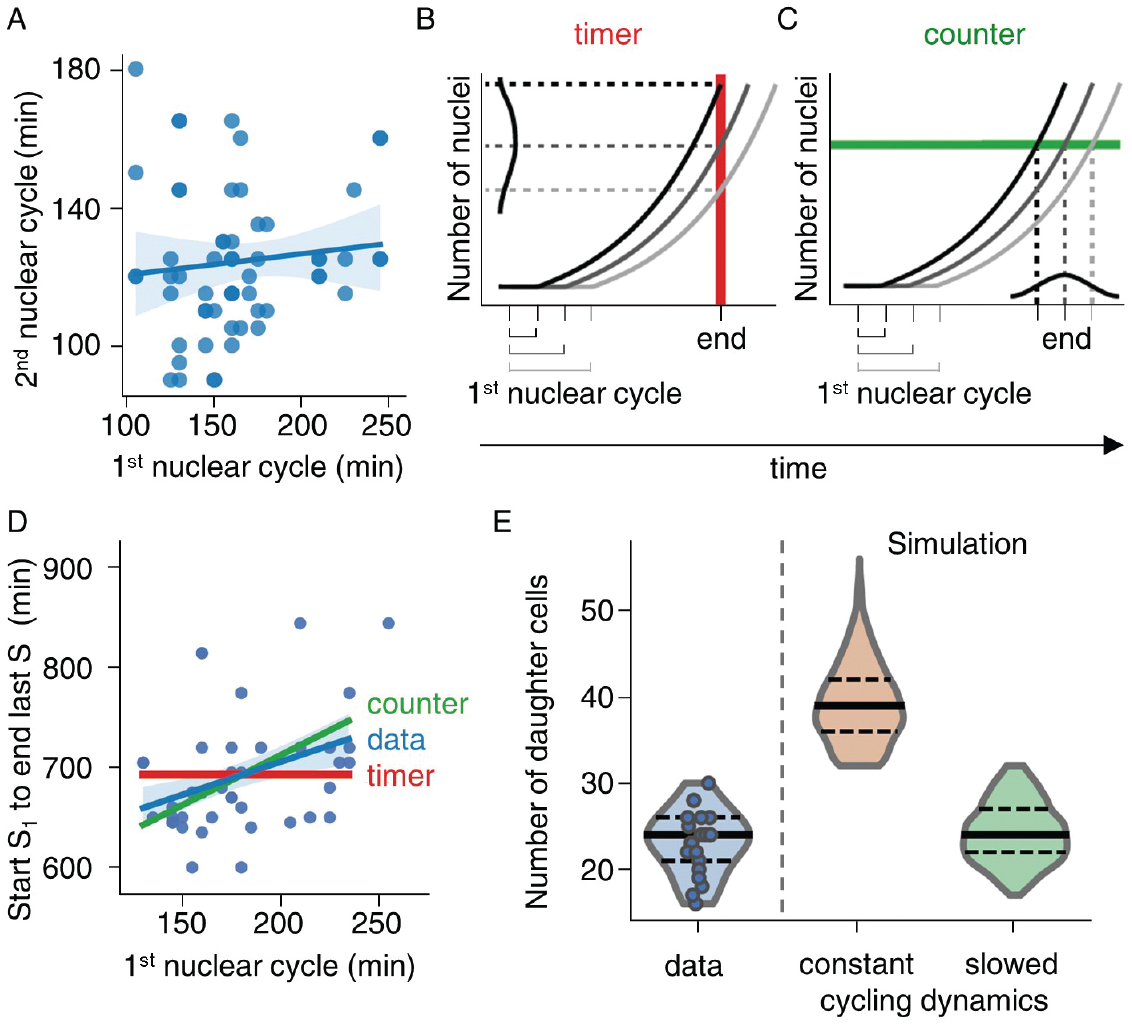
Simulation of *P. falciparum* proliferation predicts slowing nuclear cycle dynamics. (A) Time-lapse imaging of the first and second nuclear cycles showed no correlation; solid line, linear regression; band, bootstrapped 95% confidence interval; Spearman’s *ρ* = 0.14, *p* = 0.28. (B, C) Schematic illustrating how the duration of the first nuclear cycle effects the time needed to complete nuclear multiplication (defined as time from start S_1_ to end of last S-phase, i.e., last nuclear accumulation of PCNA1::GFP) in case of a timer or counter mechanism. (B) Timer mechanism with a set total time (red line) predicts no correlation between duration of first nuclear cycle and total time needed. (C) Counter mechanism with a set total number of nuclei (green line) predicts a positive correlation between the duration of first nuclear cycle and total time needed. (D) Time-lapse imaging data supports a counter mechanism (*p* = 0.077) and contradicts a timer mechanism (*p* = 0.00076); solid line, linear regression; band, bootstrapped 95% confidence interval. (E) Mathematical model with slowing nuclear cycling dynamics (17% per generation) fitted the experimental data best; solid lines, median; dashed lines, quartiles.

Because independent branching leads to very diverse trees, it is important to consider by what mechanism nuclear multiplication is stopped to achieve a well-defined endpoint that allows for cellularization. We examined two qualitatively different mechanisms, which have been proposed in the context of cell proliferation (17). A timer mechanism posits that a system is set to stop growth after an independently determined time period. This predicts that the overall duration from the start of first S-phase to end of last S-phase (i.e., end of last detected PCNA1 nuclear accumulation) (Fig. S6D) is independent of the length of the first nuclear cycle (i.e., start S_1_ to start S_2_) (Fig. 3B). Conversely, a counter mechanism stops multiplication after a certain system size (e.g., number of nuclei) has been reached, independent of the time needed. This predicts that the duration of the first cycle should correlate positively with the overall duration, as, e.g., a slow first cycle would increase the total time needed to produce a desired number of nuclei (Fig. 3C). Comparing the duration of first nuclear cycles to the overall duration showed a positive correlation (Fig. 3D and Fig. S6E–G), which is incompatible with a timer mechanism but consistent with a counter mechanism. We therefore adopted the counter mechanism and adjusted the simulation to reproduce the measured distribution of times from onset of S_1_ to the end of last S-phase (Fig. S6H, I).

To test the model, we asked how many progeny it generated overall and compared this to our experimentally determined values (Fig. 3E). When cycling dynamics were kept unchanged from nuclear cycle two onwards, simulations overestimated the total progeny. Gradually slowing nuclear cycles by 17% per generation after cycle two fitted our data best and recovered both the average and distribution in the number of progeny (Fig. 3E and Fig. S6H, I). As nuclear multiplication proceeds, the number of nuclei sharing the same cellular resources increases. Additionally, the probability that several nuclei are in the same stage of the nuclear cycle increases. Thus, the incremental decrease in nuclear cycling speed could be explained by a factor that becomes increasingly limiting as nuclei multiply. If a limiting resource needed for DNA replication was shared between nuclei, then simultaneously replicating nuclei should experience a stronger limitation than nuclei that replicated their DNA sequentially. To test this prediction, we compared pairs of nuclei were S-phases were either none, partial or completely overlapping (Fig. 4A, B). While the intervals between S-phases did not change, the duration of S-phases increased slightly from non-overlapping to partially overlapping S-phases, and strongly from partially overlapping to completely overlapping S-phases (Fig. 4C and Fig. S7A–D). This increase supports the notion that the speed of DNA replication is affected by a limiting factor. Testing if the increased duration of S-phases translated into a longer nuclear cycle, we found that partially overlapping S-phases had no effect. By contrast, a sizeable fraction of sister nuclei with synchronous S-phases displayed an increased nuclear cycle duration (Fig. 4D and Fig. S7E), suggesting that delays caused by synchronous S-phases cannot be fully compensated.

**Fig. 4.**
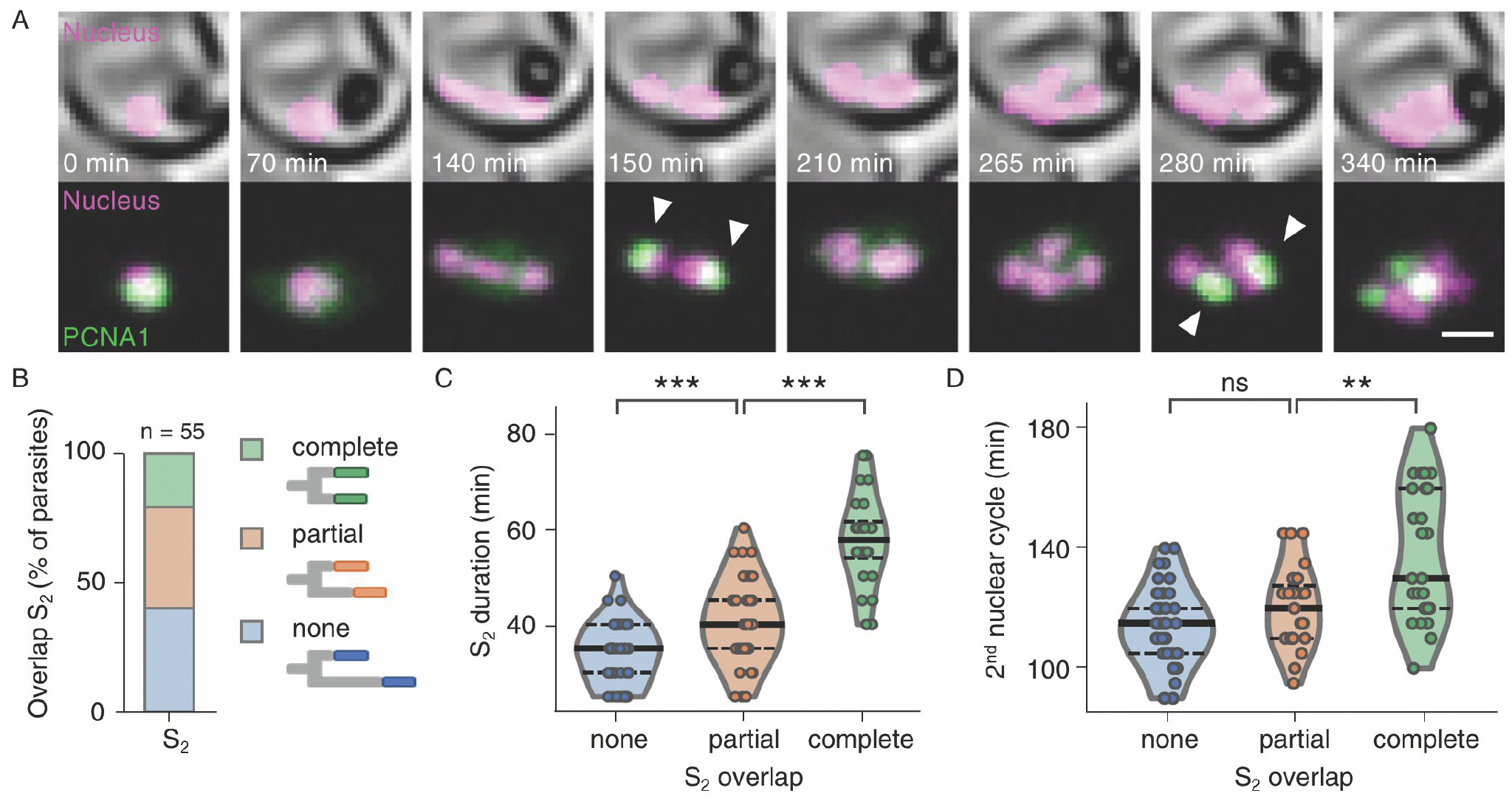
Asynchronous nuclear cycles enable rapid parasite proliferation. (A) Time-lapse microscopy of a parasite with synchronous DNA replication events (arrowheads); bar, 2 µm; movie S6. (B) Fraction of parasites with completely-, partially-, and non-overlapping S_2,1_ and S_2,2_. (C) Durations of S-phases increased with the degree of temporal overlap; solid lines, median; dashed lines, quartiles; two-sided Mann–Whitney *U* test, none and partial, *p* = 0.00027; partial and full, *p* = 5.91e^−08^. (D) Nuclear cycles containing synchronous S-phases were longer; solid lines, median; dashed lines, quartiles; two-sided Mann–Whitney *U* test, none and partial, *p* = 0.07; partial and full, *p* = 0.0067.

## Conclusions

Our results show that *P. falciparum* proliferates through alternating, consecutive rounds of DNA replication and nuclear division. Although nuclei reside in a shared cytoplasm, DNA replications and nuclear divisions occurred asynchronously. Heterogeneous and transient accumulation of PCNA1 indicates that DNA replication is regulated at the level of individual nuclei. The speed of DNA replication correlated with the degree of asynchrony, implying the existence of a limiting factor. This factor may exert an evolutionary pressure, which selects for asynchronous nuclear cycles. Our findings support a model where asynchrony enables blood-stage parasites to balance available resources with rapid proliferation, which is the main driver of virulence and critical for life cycle progression.

## Acknowledgments

We thank E. Egan for critically reading the manuscript and G. Lukinavičius and J. Bucevičius for providing the 5-SiR-Hoechst dye. We thank the Infectious Diseases Imaging Platform (IDIP) at the Center for Integrative Infectious Disease Research, Heidelberg, Germany, for the microscopy support and the Electron Microscopy Core Facilities at the European Molecular Biology Laboratory and Heidelberg University, Heidelberg, Germany. The *Plasmodium* data base PlasmoDB facilitated this work. This work was funded by the Deutsche Forschungsgemeinschaft (DFG, German Research Foundation) through project number 240245660 – SFB 1129 (C.F., T.H., F.F., U.S.S. and M.G.), and project number 349355339 (J.G.); the Baden-Württemberg Foundation (ref: 1.16101.17; M.G.); the Fundação para a Ciência e Tecnologia (FCT, Portugal) - PD/BD/128002/2016 (M.M.); P.B. was supported by the Research Training Group Mathematical Modeling for the Quantitative Biosciences.

## Author contributions

Conceptualization, S.K., P.B., N.B. U.S.S., and M.G.; Methodology, S.K., P.B., N.B., and M.G.; Software, S.K., P.B., and N.B.; Investigation, S.K., P.B., J.K., M.M., C.F., V.S., D.K., and M.C.; Formal Analysis, S.K., J.K., A.K., and P.B.; Resources, V.L., and J.G., Writing – Original Draft, S.K., P.B. N.B., and M.G.; Writing – Review & Editing, N.B., F.F., J.G., U.S.S., and M.G.; Funding Acquisition, T.H., F.F., U.S.S., and M.G.; Supervision, F.F., T.H., U.S.S., and M.G.

## Data and materials availability

All data, code, and, materials used in this analysis that are not included in the main text or in the supplementary materials are available upon request for purposes of reproducing or extending the analysis.

## Competing interests

Authors declare no competing interests

## Notes

### Competing Interest Statement

The authors have declared no competing interest.

