## Supplementary materials for "Asynchronous nuclear cycles in multinucleated *Plasmodium falciparum* enable rapid proliferation"

### Materials and Methods

***P. falciparum* cell culture.** The *P. falciparum* strain 3D7 is laboratory-adapted and derived from the isolate NF54 by limiting dilution. NF54 originated from a case of airport malaria in the Netherlands (18). *P. falciparum* 3D7 parasites were routinely cultured in human O+ peripheral blood erythrocytes in Roswell Park Memorial Institute (RPMI 1640 GlutaMAX, gibco, Thermo Fisher Scientific) medium supplemented with 0.5% albumax (gibco, Thermo Fisher Scientific), 0.2 mM hypoxanthine (c.c.pro GmbH), 25 mM HEPES (Sigma-Aldrich, Merck) pH 7.3, and 12.5 µg/ml gentamycin-sulphate (Carl Roth) (complete RPMI) at a 4% haematocrit and a temperature of 37 °C in 90% relative humidity, 3% CO2 and 5% oxygen (19). If needed, parasites were synchronized by incubating 400 µl packed erythrocytes with 10 ml of prewarmed 5% sorbitol solution (w/v) for 10 min at 37 °C (20). Parasites were washed once with complete RPMI before return to culture. Parasitemia was counted by using either a Zeiss Axiostar plus (Carl Zeiss Microscopy GmbH) or Nikon eclipse E100 (Nikon Corporation) with 100x oil immersion objective on thin blood smears fixed with 100% methanol (Honeywell) for ten seconds and stained with Hemacolor rapid staining of blood smears solution (Sigma-Aldrich, Merck). For genomic DNA extraction, parasite DNA was isolated from 200 µl washed erythrocyte pellet with a parasitemia of 1-3% by use of the DNeasy blood & tissue kit (Qiagen).

**Generation of transgenic *P. falciparum*.** All primer and plasmid sequences are available upon request. The plasmid for visualization of parasite nucleoplasm was generated by restriction digest mediated removal of the FKBP-rapamycin binding domain (FRB) sequence from the plasmid p3xNLS-FRB-mCherry-hsp86-BSD gifted by Tobias Spielmann (21). The resulting plasmid p3xNLS-mCherry-hsp86-BSD expresses mCherry coupled to 3 nuclear localization sequences (NLS) under the constitutive hsp86 promoter. Furthermore, the plasmid contains a blasticidin (BSD) resistance cassette.

PCNA1::GFP was expressed from the plasmid pARL_PCNA1-GS-eGFP under the control of the *P. falciparum* CRT promoter. Additionally, it contains a WR99210 resistance cassette (22). Plasmids were amplified by transformation into 50 µl competent 5-alpha F'Iq Competent *E. coli* cells (New England Biolabs GmbH), overnight culture at 37 °C and extraction via GenElute HP Plasmid Miniprep Kit (Sigma-Aldrich, Merck) or NucleoBond Xtra Midi kit (Macherey-Nagel) depending on required purity and yield. Monoclonal colonies were verified by Sanger sequencing (Eurofins Genomics).

Transgenic parasites were created by electroporation (310 kV, 950 µF; Gene Pulser II, Bio-Rad) of 50-100 µg of purified DNA resuspended in 30 µl TE buffer (10 mM Tris pH 8.0, 1 mM EDTA pH 8.0) and 370 µl cytomix (120 mM KCl, 0.15 M $CaCl_2$, 2 mM EGTA, 5 mM $MgCl_2$, 10 mM $K_2HPO_4$/$KH_2PO_4$, 25 mM HEPES, pH 7.6) into 200 µl packed erythrocytes of a ring-stage 3D7 culture with a parasitemia of at least 4% (23). Presence of episomes was maintained by continuous selection with blasticidin (InvivoGen) at 5 µg/ml and/or WR99210 (Jacobus Pharmaceutical Company) at 2.5 µM, respectively.

Endogenous tagging of PCNA1 with GFP was attempted by CRISPR/Cas9-mediated homology repair. The plasmid pDC2-cam-coCas9-U6-hDHFR was a kind gift from Marcus Lee (24). Briefly, this plasmid expresses a guide RNA (gRNA) sequence from a U6 cassette together with a the Cas9 nuclease under the *P. falciparum* calmodulin promoter as well as a WR99210 resistance cassette for selection in *P. falciparum*. gRNA binding sites were selected using the Protospacer Workbench software (25). The selection of gRNA sequences was based on distance to the C-terminus of PCNA1 as well as the activity prediction scores (26). gRNAs were ordered as unmodified oligonucleotides (Thermo Fisher Scientific), annealed and phosphorylated before ligation in the pDC2-cam-coCas9-U6-hDHFR plasmid linearized by BbsI restriction digest (New England Biolabs GmbH). Template regions for double-crossover homology repair were cloned into the same plasmid backbone. The final plasmids were assembled via Gibson assembly (NEBuilder HiFi DNA Assembly, New England Biolabs GmbH) using the PCR-amplified homology- and GFP sequences and the plasmid containing the selected guides, which was linearized by EcoRI and AatII restriction digest (New England Biolabs GmbH).

Transfection was carried out as described above, but selection with WR99210 was discontinued 12 days after transfection. Presence of GFP fluorescence was investigated in live parasites under a cover slip on a glass slide using a 60x oil immersion objective on a Zeiss Axio Observer (Carl Zeiss Microscopy GmbH) equipped with a Prime BSI sCMOS camera (Teledyne Photometrics). Diagnostic PCR was carried out on isolated genomic DNA from the transfected pool.

**Sample preparation for correlative light and electron tomography and serial section tomography.** High pressure freezing: Red blood cells infected with *P. falciparum* 3D7 expressing 3xNLS-mCherry and PCNA1::GFP in multinucleated stages were purified via varioMACS (Miltenyi Biotec GmbH) magnetic purification, washed in PBS (gibco, Thermo Fisher Scientific) and resuspended in incomplete RPMI (0.2 mM hypoxanthine, 25 mM HEPES pH 7.3, 12.5 µg/ml gentamicin). The pellet was pipetted into 0.2 mm deep aluminum carriers (Engineering Office M. Wohlwend GmbH) and cryo-immobilized by high pressure freezing using Leica EM ICE (Leica Microsystems). Freeze substitution was done using a freeze substitution device EM-AFS2 (Leica Microsystems) and the freeze substitution solution contained 0.3% (w/v) uranyl acetate dissolved in anhydrous acetone and the samples were substituted at -90 °C for 14 h. The temperature was then increased at a rate of 5 °C/h to -45 °C followed by 2 h incubation at -45 °C. The samples were rinsed with ethanol for 1 h, followed by LR Gold (London Resin Company) infiltration at -25 °C in 2 h steps with 25%, 50% and 75% LR Gold in ethanol. The samples were then left in 100% LR Gold for 12 h and 4 h before the onset of polymerization. UV polymerization was applied for 24 h at -25 °C and the temperature was increased to 20 °C at a rate of 5 °C per hour. The samples were left exposed to UV at room temperature for 24 h.

**Correlative light and electron microscopy.** Sectioning was done on a Leica UC6 ultramicrotome (Leica Microsystems) and sections were collected on formvar coated finder grids (Plano). 200 nm thick sections on grids were placed in PBS and sandwiched between two cover glasses and fluorescence signal was imaged on a Zeiss observer Z1 fluorescence microscope (Zeiss). The identical cells were imaged on a JEOL JEM-1400 electron microscope (JEOL) operating at 80 kV and equipped with a 4K TemCam F416 camera (Tietz Video and Image Processing Systems GmBH). In the final step, the correlation of fluorescence and electron micrographs, based on morphological features, was done using the eC-CLEM software (27).

**Serial section tomography.** Electron tomography: The resin block was sectioned in 200 nm sections using a Leica UC7 ultramicrotome (Leica Microsystems). Ribbons of serial sections were placed onto formvar-coated slot copper grids and imaged with a Tecnai F30 transmission electron microscope (FEI) operating at 300 keV. Tilt-series were recorded at the angle ranging from -60° to 60° with 2° increments using SerialEM software (28). Digital images were recorded on a fast Gatan OneView 4k camera with a nominal magnification of 12,000. Image processing and analysis: 3D tomograms were generated from the tilt-series using weighted back projection and combined into serial tomograms using IMOD software package (29). Schizont nuclei were segmented using AMIRA software (Thermo Fischer Scientific). Minimal distance between adjacent nuclei was measured with the line tool in ImageJ by manually connecting the edge of the nucleoplasm of neighboring nuclei.

**Live *P. falciparum* labelling and imaging.** For live-cell imaging parasites were seeded on sterile glass bottom 35 mm µ- or 8 well-dishes (ibidi GmbH) as described by Grüring and Spielmann with minor modifications (30). Briefly, the bottom of the dish was coated with 5 mg/ml concanavalin A (Sigma-Aldrich) and rinsed with PBS. Resuspended parasite culture was washed twice with incomplete RPMI and left to settle on the dish for 10 min at 37 °C before unattached cells were washed off using incomplete RPMI until a single-cell layer remained. Cells were left to recover at standard culturing conditions in complete RPMI for at least 8 h before media was exchanged to phenol red-free complete RPMI imaging medium (RPMI 1640 L-Glutamine (PAN-Biotech), 0.5% albumax, 0.2 mM hypoxanthine, 25 mM HEPES pH 7.3, 12.5 µg/ml gentamicin) that had been equilibrated to incubator gas conditions for at least 6 h. Dishes were completely filled, closed and sealed tightly using parafilm before imaging was started. For staining of parasite DNA, phenol red-free complete RPMI imaging medium was supplemented with 5-SiR-Hoechst kindly provided by Gražvydas Lukinavičius und Jonas Bucevičius (13). 5-SiR-Hoechst was added at 250 nM in imaging medium to parasites 2 h prior to imaging.

Routine long-time live cell imaging was carried on a PerkinElmer UltraVIEW VoX microscope equipped with Yokogawa CSU-X1 spinning disk head and Nikon TiE microscope body. An Apo TIRF 60x/1.49 N.A. oil immersion objective and Hamamatsu C9100-23B EM-CCD camera were used. Live-cell imaging was performed at 36.5˚C. Image were acquired at multiple positions using an automated stage and the Perfect Focus System (PFS) for focus stabilization with a time-resolution of 5 min/stack. Multichannel images were acquired sequentially using solid state lasers with excitation at 488 nm and 561 nm and matching emission filters in addition to differential interference contrast (DIC) images; 8 µm stacks were acquired with a z-spacing of 500 nm.

Live-cell DNA quantification in parasites containing one nucleus was performed at 37 °C on a PerkinElmer UltraVIEW VoX microscope equipped with Yokogawa CSU-X1 spinning disk head and Nikon TiE microscope body. A Plan Apo VC 100x/1.4 N.A. oil immersion objective and Hamamatsu C9100-23B EM-CCD camera were used in combination with an additional zoom of 1.5x. Live-cell imaging was performed at 36.5 °C. Image were acquired at multiple positions using an automated stage and the Perfect Focus System (PFS) for focus stabilization with a time-resolution of 5 min/stack. Multichannel images were acquired sequentially using solid state lasers with excitation at 488 nm, 561 nm and 640 nm and matching emission filters in addition to DIC images; 6 µm stacks were acquired with a z-spacing of 500 nm.

Additional live-cell DNA quantification in parasites with one or two nuclei was done on a point laser scanning Leica SP8 microscope using HC PL APO CS2 63x/1.4 N.A. oil immersion objective. Image were acquired at multiple positions using an automated stage and the Adaptive Focus Control (AFC) for focus stabilization with a time-resolution of 5 min/stack. Multichannel images were acquired sequentially using a PMT detector with a gain of 800-1000 V, laser excitation at 561 nm and a spectral detection window set between 570 nm and 620 nm for mCherry imaging and using a HyD detector in the standard mode with laser excitation at 633 nm and spectral detection window set between 640 and 700 nm for 5-SiR-Hoechst. In parasites with one nucleus, additional images were acquired using a HyD detector in the standard mode with laser excitation at 488 nm and spectral detection window set between 490 and 550 nm for GFP imaging. Brightfield images were obtained from a transmitted light PMT detector. A 2.25-4.5 µm stack was acquired with a sampling of 60-100 nm in xy axis and 0.75-1 µm in z axis with scanning speed of 400-700 Hz.

For DNA quantification in fixed cells, *P. falciparum* infected erythrocytes were seeded in 8 well-dishes (ibidi GmbH) in the same manner as for live-cell imaging. After a minimum of 4h recovery time in complete RPMI, cells were fixed using prewarmed 4% paraformaldehyde (Electron Microscopy Sciences) in PBS for 20 min at 37 °C, before washing with PBS. For staining of DNA, 5-SiR-Hoechst at a concentration of 250 nM was added to PBS 30 minutes prior to imaging.

Point laser scanning confocal microscopy was performed on a Leica SP8 microscope using a HC PL APO CS2 63x/1.4 N.A. oil immersion objective. Images were acquired using HyD detectors in the standard mode with the laser excitation at 633 nm and spectral detection window set between 640 and 700 nm for 5-SiR-Hoechst imaging. Brightfield images were obtained from a transmitted light PMT detector. A 6 µm stack was acquired with a sampling of 29 nm in xy axis and 132 nm in z axis with scanning speed of 400 Hz. Subsequently, images were processed with the Lightning algorithm in the adaptive mode using default settings.

**Image analysis.** For Image analysis was done using the Fiji distribution of ImageJ (31). Quantitative results were exported as comma-separated value (csv) files, analyzed and plotted using Microsoft Excel, GraphPad Prism version 5.0.0 for Windows or using python with matplotlib 3.2.2, numpy 1.19.2, pandas 1.2.0 and scipy 1.6.0. DNA quantification in fixed cells: Nuclei were counted by three independent researchers in Lightning algorithm-processed 3D stacks. For DNA intensity measurements, 5-SiR-Hoechst signal in the total area occupied by nuclei was measured in individual parasites. Briefly, a threshold (300, 65536) was applied to Lightning algorithm-processed 3D stacks and the resulting mask used to measure raw integrated density of the 5-SiR-Hoechst channel within segmented areas of unprocessed images. Signal intensities were normalized to the mean 5-SiR-Hoechst signal intensity in nuclei of ring stage parasites.

Live-cell DNA quantification in parasites with two nuclei: For quantification of DNA in live-cell microscopy, 5-SiR-Hoechst signal intensities in individual nuclei, segmented via the mCherry signal, were measured over time. Areas with individual nuclei were manually separated. A median filter with a radius of two pixel was applied to the mCherry signal of individual nuclei and a threshold (20, 255-35, 255) depending on the fluorescence intensity of mCherry in the cell, was applied to create masks of individual nuclei. 5-SiR-Hoechst signal intensity was then determined in the individual nuclei via measurement of raw integrated density within nuclear masks. Signal intensities in individual nuclei were normalized to the first timepoint of the time course or to the timepoint in which individual nuclei were first observed.

Live-cell DNA quantification in parasites with one nucleus: For parasites that had one nucleus and expressed PCNA1::GFP, the 5-SiR-Hoechst signal intensity in the nucleus was measured over time and correlated to the maximal PCNA1::GFP signal. Data acquired on the Perkin Elmer spinning disk microscope was processed as follows: background values for each channel were determined by the mean signal in an area not containing a parasite. The mCherry signal was used to determine the nucleus area. A background subtraction of 2020 as well as a median filter with a radius of one pixel was applied to the mCherry signal of individual cells, followed by a threshold (170, 65536) to create masks of individual nuclei. 5-SiR-Hoechst signal intensity was then determined in the individual nuclei via measurement of raw integrated density after background subtraction of 2300. Signal intensities were normalized to the average of the first ten values of the time course or to the average of all available values from the start of the time course to the start of nuclear PCNA1::GFP accumulation if both timepoints were less than ten timepoints apart from each other. In addition, maximal GFP signal intensity values were measured from an average intensity z-projection of the GFP channel and normalized to minimal and maximal values for each cell. Data from individual cells were aligned to the timepoint at which the highest maximal GFP signal intensity occurred.

Data acquired on the Leica Sp8 confocal laser-scanning microscope was processed as follows: The mCherry signal was used to determine the nucleus area. A median filter with a radius of one pixel was applied to the mCherry signal of individual cells, followed by a threshold (15, 255) to create masks of individual nuclei. 5-SiR-Hoechst signal intensity was then determined in the individual nuclei via measurement of raw integrated density after background subtraction of 10. The area of the nucleus was determined via measurement of the area of the mCherry mask and 5-SiR-Hoechst signal intensity was divided by the nuclear area to avoid total signal variations due to changing z-focus during imaging. Signal intensities per area were normalized to the average of the first ten values of the time course or to the average of all available values from the start of the time course to the start of nuclear PCNA1::GFP accumulation if both timepoint were less than ten timepoints apart from each other. In addition, maximal GFP signal intensity values were measured from an average intensity z-projection of the GFP channel and normalized to minimal and maximal values for each cell. Data from individual cells were aligned to the timepoint at which the highest maximal GFP signal intensity occurred.

Analysis of long-term live-cell imaging: Cells were manually evaluated for parasite survival and health and grouped into different categories depending on which events of schizogony were clearly observable, i.e., onset of S-phase, number of nuclei at start of imaging, parasite egress, and start of first S-phase and parasite egress in the same cell. Individual parasites were manually analyzed for start and end of S-phase using nuclear PCNA1::GFP accumulation as proxy, and for timepoints of nuclear division, i.e., earliest observable occurrence of completely separate nuclei. Determination of GFP fluorescence intensity before first nuclear PCNA1::GFP accumulation: Timepoints for determination of GFP fluorescence intensity before start of nuclear PCNA1::GFP accumulation were manually selected. Background intensity was measured by determining maximum fluorescence in an area not occupied by the parasite and subtracted from the GFP channel. Subsequently, the area containing the parasite and raw integrated density of the GFP channel within this area was measured. Analysis of fluorescence intensity over time: To analyze fluorescence levels over time, representative parasites were chosen where both onset of first S-phase and egress were clearly visible. Raw integrated density was measured in the mCherry and GFP channel of average intensity z-projections and normalized to maximal and minimal values per cell. Data from individual cells were aligned by the timepoint of parasite egress.

Visualization of nuclear PCNA1::GFP accumulation over time: To visualize occurrence of PCNA1::GFP accumulation over time, normalized maximal GFP pixel intensity values over time were plotted as a heatmap. Maximal GFP pixel values per timepoint were measured from average intensity z-projections and normalized to maximal and minimal values per cell. Cells were aligned by occurrence of first PCNA1::GFP nuclear accumulation and ordered by occurrence of second PCNA1::GFP accumulation. The heatmap was created in ImageJ by re-normalizing values to the 8-bit value range (0-255) importing values as text image.

Determination of GFP fluorescence within nucleus and cytoplasm: Four representative parasites were chosen to analyze levels of PCNA1::GFP in the nucleus and in the cytoplasm over time. Fifty consecutive timepoints were chosen manually that showed parasites with one nucleus and an increasing and decreasing nuclear PCNA1::GFP signal. Background was determined for each parasite by measuring mean and maximum GFP values in an area not containing a parasite and set at $x_{\text{background}} = x_{\text{mean}} + \frac{x_{\text{max}} - x_{\text{mean}}}{2}$. $x_{\text{background}}$ was subtracted from the GFP channel in each plane of the 3D stack at every timepoint. The area of the nucleus was identified via automatic thresholding in the mCherry channel to create a nuclear mask. Subsequently, raw integrated density of the GFP channel was determined per timepoint in the total parasite area as well as in the nucleus. Values for the cytoplasm were calculated by subtracting values for the nucleus from the values of the total parasite. Data from individual cells were aligned by occurrence of highest nuclear PCNA1::GFP signal.

**Branching process models for nuclear multiplication.** General setup: To systematically explore the temporal evolution of nuclear multiplication during schizogony, we construct a mathematical model of nuclear multiplication as a branching process. Within this process, each nuclear cycle is divided into two phases, S and D (Fig. S6A). During S-phase the genome is replicated within a single nucleus. The S-phase is followed by two parallel D-phases, one for each daughter nucleus, which comprise the time between the end of DNA replication in the mother and the beginning of the next round of replication in the daughters. Thus, the D-phase in the i-th nuclear cycle corresponds to the $SD_i$ and $DS_{i+1}$ phases as defined in the main text. Each D-phase spans one nuclear division whose exact time is not resolved in the model; instead we opted for the minimal model that captures individual replications. For simplicity, we model the phase durations as independent random times. This is accurate for adjacent S- and D-phases, as well as D- and next-cycle D-phases, which are not significantly correlated in our data. We also neglect the correlation of S- and subsequent S-phases (Fig. S6B) and between S- and D-phases in sister nuclei. While these were significantly positive in our data, including them had little effect on the predictions of our model, and we therefore do not present these details here. In sum, each (S- or D-) phase duration is independently sampled from a Gamma distribution with density, $f(x;\, \alpha, \beta) = \beta^{\alpha} x^{\alpha-1} e^{-\beta x}/\Gamma(\alpha)$ parameterized by experimental data. Specifically, realizations of the nuclear branching process are generated as follows. At time $t = 0$ a single nucleus enters S-phase ($S_1$), with a duration drawn from the corresponding Gamma distribution (Tab. S1). The two subsequent D-phases ($D_{2,1}$ and $D_{2,2}$) are then sampled independently from the corresponding distribution $D_2$ (Tab. S1). For nuclei of all following generations, this procedure is repeated, the only difference being that the durations for S- and D-phases are somewhat faster (Tab. S1). In particular, as cycle phases were measured directly only for the first and second nuclear cycles, we initially adopt the hypothesis that all subsequent nuclear cycles share the statistics of the second cycle.

Stopping the simulation: The branching process ends when a stopping criterion is met. Arguably the simplest criteria are stopping based on elapsed total time, called timer mechanism, or stopping based on total size of the system, called sizer mechanism (17). The timer mechanism predicts no correlation between the duration of the first nuclear cycle and the total time until nuclear multiplication stops, but our data showed a positive correlation (Fig. 3D). We therefore opted for the sizer mechanism, which generically produces a positive correlation between the completion of an initial event and the overall process. Concretely, we implement the sizer mechanism based on counting of nuclei, and therefore call it a counter mechanism in the following and throughout the main text. After reaching a given number of nuclei $n_{\text{stop}}$, all running DNA replications are completed but no further replications may be initiated. Thus, the final number of nuclei $n$ is given by $n = n_{\text{stop}} + n_{\text{running}}$, where $n_{\text{running}}$ is the number of running replications at the time $n_{\text{stop}}$ nuclei are reached.
To parameterize the stopping condition of our branching process, we adjusted $n_{\text{stop}}$ to reproduce the total duration of replication. By simulating 10,000 schizogonies with $n_{\text{stop}} = 32$, both median and interquartile range of the measured total duration of replication were recovered (Fig. S6H). Furthermore, the counter mechanism reproduced the positive correlation between the first nuclear cycle and the overall time of replication (Fig. S6I).

Predictions for the total merozoite count: With the cycling parameters above and $n_{\text{stop}} = 32$ the model predicts $n = 39\ (36, 42)$ [median, (1st, 3rd quartiles)] merozoites in total, whereas $24\ (21, 26)$ were measured (Fig. 3E). As we observed no loss of nuclei that participated in multiplication by time lapse microscopy, this implies that the simulated third and later nuclear cycles are too quick on average. In particular, the assumption of unchanged cycling statistics from the second cycle onwards needed to be revised: cycling speeds up from cycle one to two but then slows down again as schizogony continues. To estimate the slowdown of nuclear cycling after the second cycle, we modified our previous branching model in a parsimonious way, by introducing one new parameter, the retardation factor $\gamma$. In the modified model, starting from the third nuclear cycle, each S- or D-phase is prolonged by a factor of $\gamma$ per cycle. For example, the S-phase S4 in cycle $i = 4$ is sampled from a Gamma distribution with unchanged $\alpha = 12.49$ (Tab. S1) but reduced rate parameter $\beta = \beta/\gamma^{i-2} = \beta/\gamma^2$. Compared to an alternative model where all nuclear cycles $i \geq 3$ are statistically identical and slower than the second cycle, this gradual slowdown has the advantage of avoiding a step-like change in cycling speed. We then re-fitted the branching process model. 10,000 simulated schizogonies with $\gamma = 1.17$ and $n_{\text{stop}} = 17$ show that both median and interquartile range of the total duration of nuclear multiplication (Fig. S6H), the final number n of merozoites (Fig. 3E) and the positive initial-full correlation (Fig. S6I) are captured accurately by the branching process with slowdown. In particular, this model predicts that the final S-phases, which appear in nuclear cycle $i = 5$, are on average 60% longer than S2.

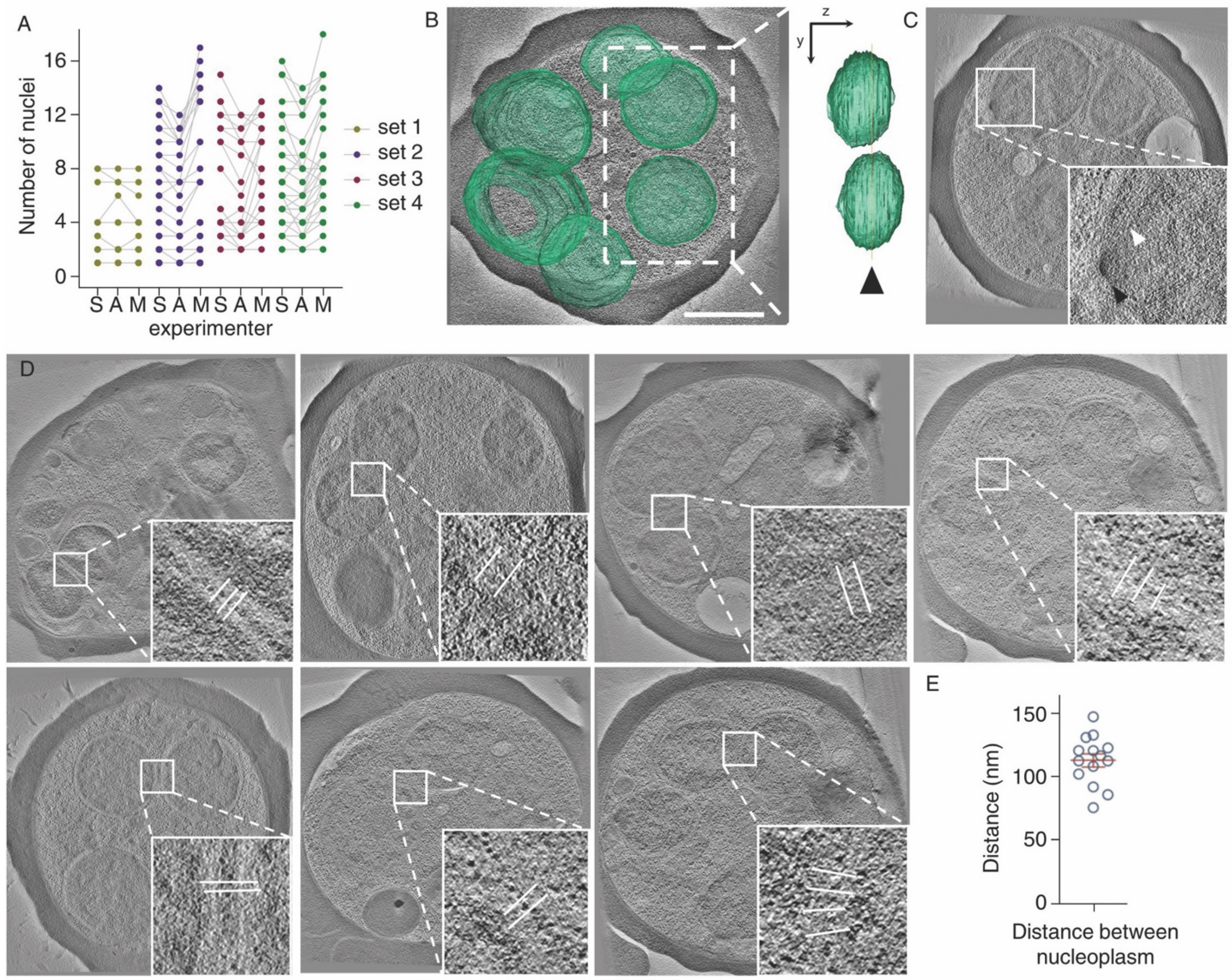


**Fig. S1. Nuclei of a *P. falciparum* schizonts are separate cellular compartments except during division** (A) Quantification of *P. falciparum* nuclei by three independents experimenters (S., A., and M.). Sets of parasites from independent biological replicates. (B) Electron tomogram slice, overlayed with the 3D-segmented inner nuclear membraned (green) and side view (90° rotation around the y axis) of nuclear volumes showed complete separation; arrowhead, indicates the shown tomogram plane; bar, 1 $\mu$m; movie S2. (C) Electron tomogram of nucleus undergoing division; white arrowhead, microtubule; black arrowhead, centriolar plaque. (D) Measurements of the minimal distance between the nucleoplasm of adjacent nuclei; white lines in inserts, quantified distances; note, ribosomes were detectable in the space between nuclei. (E) Average (± standard deviation) minimal distance between the nucleoplasm of adjacent nuclei.

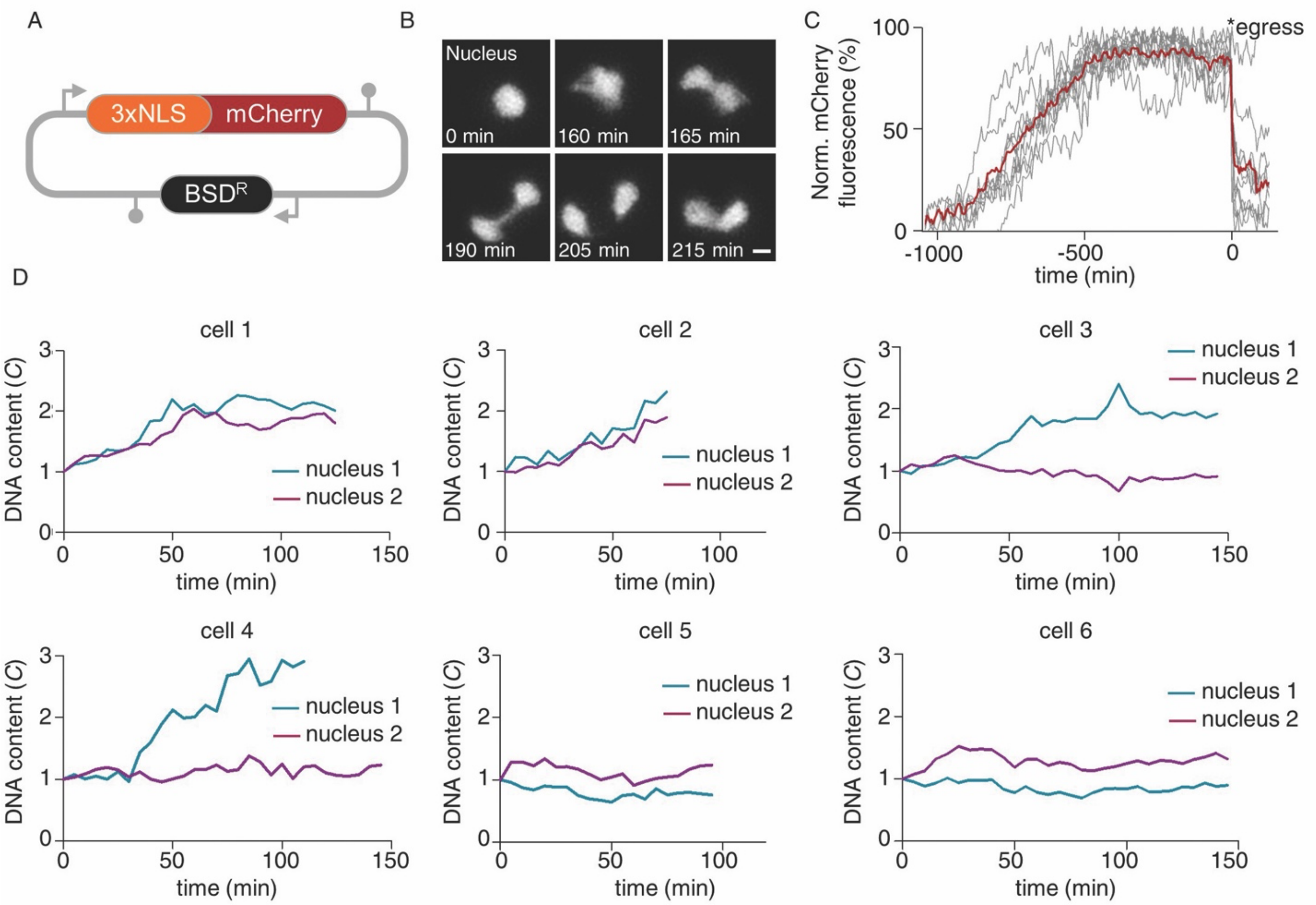


**Fig. S2. DNA replication can occur asynchronously in *P. falciparum* nuclei of the same cell.** (A) Scheme illustrating the plasmid design to generate a reporter parasite with red-fluorescent nuclei. mCherry was N-terminally fused to a triple nuclear localization signal, 3xNLS, and expressed under the control of own promoter and terminator regions. In addition, the plasmid contained a cassette expressing blasticidin-S-deaminase, conferring resistance to blasticidin S ($BSD^R$); not drawn to scale. (B) High-resolution time-lapse microscopy of a nuclear division event, nuclei express 3xNLS::mCherry; bar, 1 $\mu$m. (C) Long-term time-lapse microscopy of the 3xNLS::mCherry reporter parasite showed that the expression of mCherry is relatively constant for approximately 500 minutes before *P. falciparum* egress, i.e., daughter cell release from the host erythrocyte; data from three biological replicates; grey lines, mCherry signal of eleven individual parasites; red line, average. (D) Time-lapse microscopy of the 3xNLS::mCherry reporter parasite stained with the DNA dye 5-SiR-Hoechst and quantification of the nuclear DNA content in individual sister nuclei showed asynchronous and synchronous DNA replication events; note, the nuclei of cells 5 and 6 did not replicate during the imaging.

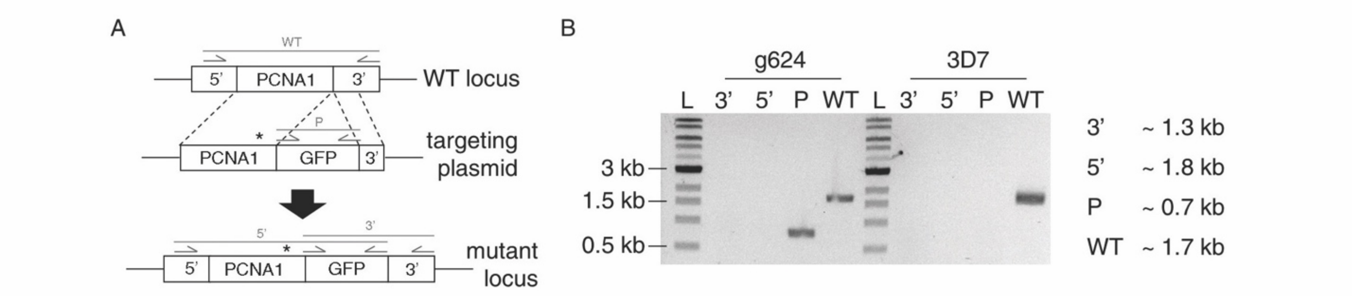


**Fig. S3. Endogenous fusion of *P. falciparum* PCNA1 with GFP via genome editing failed.** (A) Scheme illustrating the CRISPR-based genome editing strategy to fuse endogenous *P. falciparum* PCNA1 with GFP; arrows, primer binding sites; grey lines, expected PCR-amplicons of diagnostic PCR reactions; asterisk, position of mutated PAM-site; not drawn to scale. (B) Diagnostic PCR failed to detect the desired endogenous PCNA1::GFP fusion; note, endogenous tagging was attempted with two different guide RNAs; transfection with guide RNA g738 yielded non-fluorescent parasites; transfection with guide RNA g624 yielded fluorescent parasites; g624, transfectant genomic DNA; 3D7, parental wild type *P. falciparum* genomic DNA; L, 1 kb ladder; 3', 3' test PCR; 5', 5' test PCR; P, plasmid test PCR; WT, wild type test PCR.

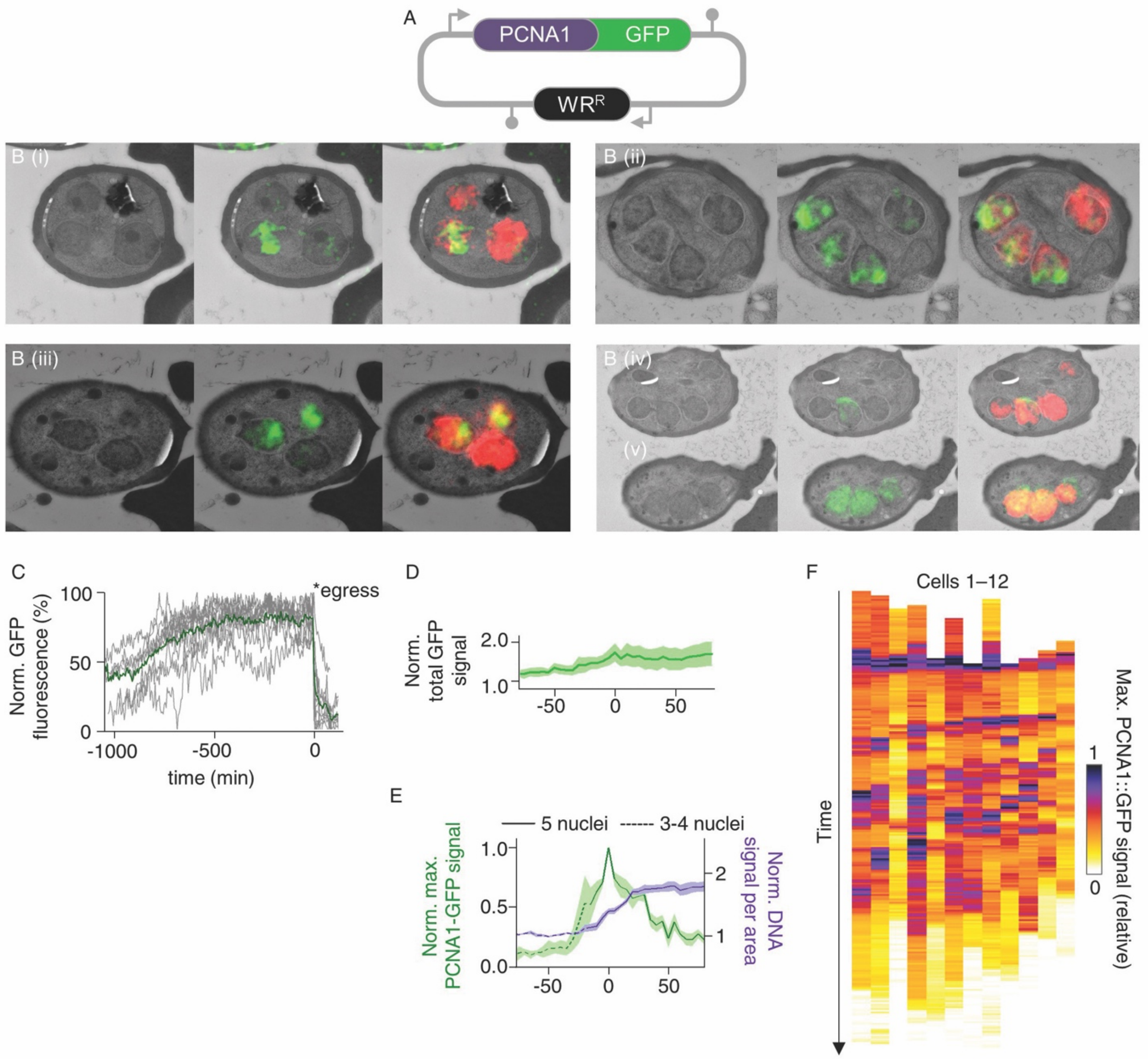


**Fig. S4. Characterization of a *P. falciparum* reporter line, episomally expressing PCNA1::GFP and 3xNLS::mCherry.** (A) Scheme illustrating the plasmid design to generate a reporter parasite for episomal expression of a *P. falciparum* PCNA1::GFP fusion in the 3xNLS::mCherry background. PCNA1::GFP was expressed under the control of own promoter and terminator regions. In addition, the plasmid contained a cassette expressing the human dihydrofolate reductase, conferring resistance to WR99120 ($WR^R$); not drawn to scale. (B) Correlative light and electron microscopy of five *P. falciparum* schizonts (i-v) showed heterogeneous accumulation of PCNA1 among nuclei; left panel, transmission electron micrograph; middle panel, transmission electron micrograph overlayed with GFP signal; right panel, transmission electron micrograph overlayed with GFP and mCherry signal. (C) Long-term time-lapse microscopy of the PCNA1::GFP, 3xNLS::mCherry reporter parasite showed that the expression of GFP is relatively constant for approximately 500 minutes before *P. falciparum* egress, i.e., daughter cell release from the host erythrocyte; data from three biological replicates; grey lines, GFP signal of eleven individual parasites; green line, average. (D) Normalized total PCNA1::GFP signal intensity over 150 minutes, corresponding to Fig. 2C; lines, average (n = 5); bands, standard deviation. (E) Nuclear PCNA1::GFP accumulation caused a peak in the maximal signal intensity, which coincided with a DNA content duplication; solid lines, average; bands, standard deviation; note, data for Fig. 2D were acquired on a Leica SP8 laser scanning confocal microscope and data for Fig. S4E were acquired on a PE spinning-disc confocal microscope. (F) Normalized maximal PCNA1::GFP-signal intensity over the course of schizogony of twelve individual parasites as proxy for nuclear accumulation of PCNA1. Heatmaps were aligned at the first peak and ordered by the time to second peak.

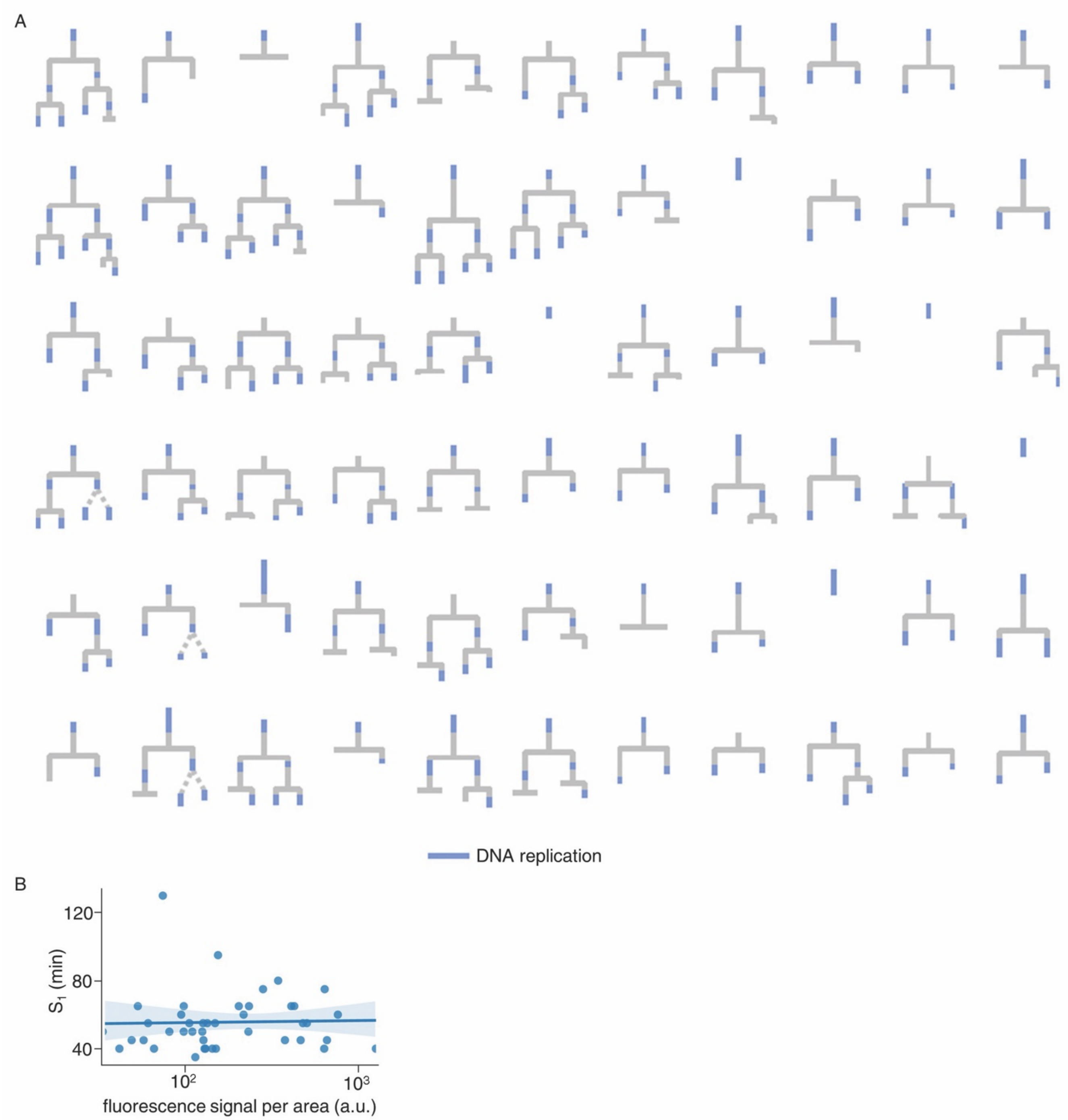


**Fig. S5. Overexpression of PCNA1::GFP allows tracing of lineage trees of *P. falciparum* nuclei and does not affect S-phase duration.** (A) Summary of all lineage trees that were analyzed for Fig. 2F–H, Fig. 3A, D, and Fig. 4B–D; dashed lines, timing of nuclear division event could not be determined with confidence; timing drawn to scale. (B) The abundance of episomally expressed PCNA1::GFP did not correlate with the duration of the first S-phase (S1); a.u., arbitrary units; solid line, linear regression; band, bootstrapped 95% confidence interval; Spearman's $\rho = 0.14$, $p = 0.37$.

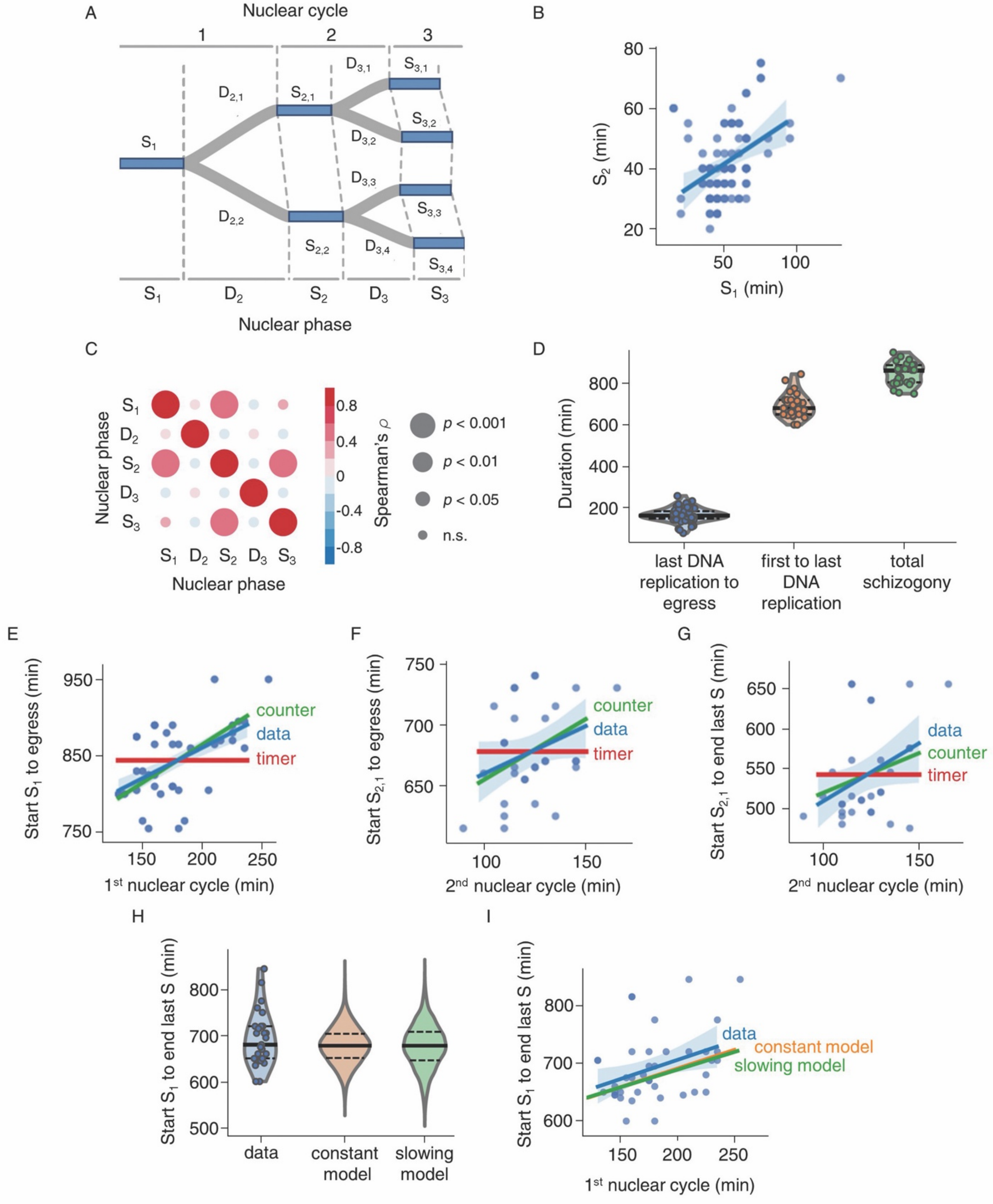


**Fig. S6.** Parametrization of a mathematical model to simulate nuclear multiplication during *P. falciparum* blood stage schizogony. (A) Scheme of the simplified two-phase branching process model. The intervals SD and DS were combined to D; generations of nuclei are indicated above the tree; dashed lines connect the onset and conclusion of S-phases within the generations. (B) Duration of the first S-phase ($S_1$) correlated with the duration of the second S-phase ($S_{2,1}$ and $S_{2,1}$); solid line, linear regression; band, bootstrapped 95% confidence interval; Spearman's $\rho = 0.44$, $p = 2.8e^{-05}$. (C) Spearman rank correlation of the phases of the two-phase tree. (D) Quantification of key parameters of *P. falciparum* schizogony via long-term time-lapse microscopy; last DNA replication was defined as last detectable nuclear accumulation of PCNA1::GFP; solid lines, median; dashed lines, quartiles. (E–G) Live-cell imaging data supports a counter mechanism; solid line, linear regression; band, bootstrapped 95% confidence interval. (E) Correlation of the duration from the start of $S_1$ to egress, i.e., parasite exit from the host erythrocyte, with the duration of the first nuclear cycle is consistent with a counter mechanism (timer, $p = 3.32e^{-07}$; counter, $p = 0.17$). (F) Correlation of the duration from the start of S2 to egress with the duration of the second nuclear cycle is consistent with a counter mechanism (timer, $p = 0.079$; counter, $p = 0.63$). (G) Correlation of the duration from the start of $S_{2,1}$ to last S-phase with the duration of the second nuclear cycle is consistent with a counter mechanism (timer, $p = 0.036$; counter, $p = 0.5$). (H) Time needed to complete nuclear multiplication. Measured data were compared to optimized solutions based on our computational model with independent nuclei and a counter mechanism (with constant cycling-speed or 17% slowdown per nuclear cycle, respectively); solid lines, median; dashed lines, quartiles. (I) The counter mechanism of our model (with constant cycling-speed or 17% slowdown per nuclear cycle, respectively) can reproduce the observed positive correlation between duration of first nuclear cycle and the duration from the start of $S_1$ to last S-phase; solid line, linear regression; band, bootstrapped 95% confidence interval.

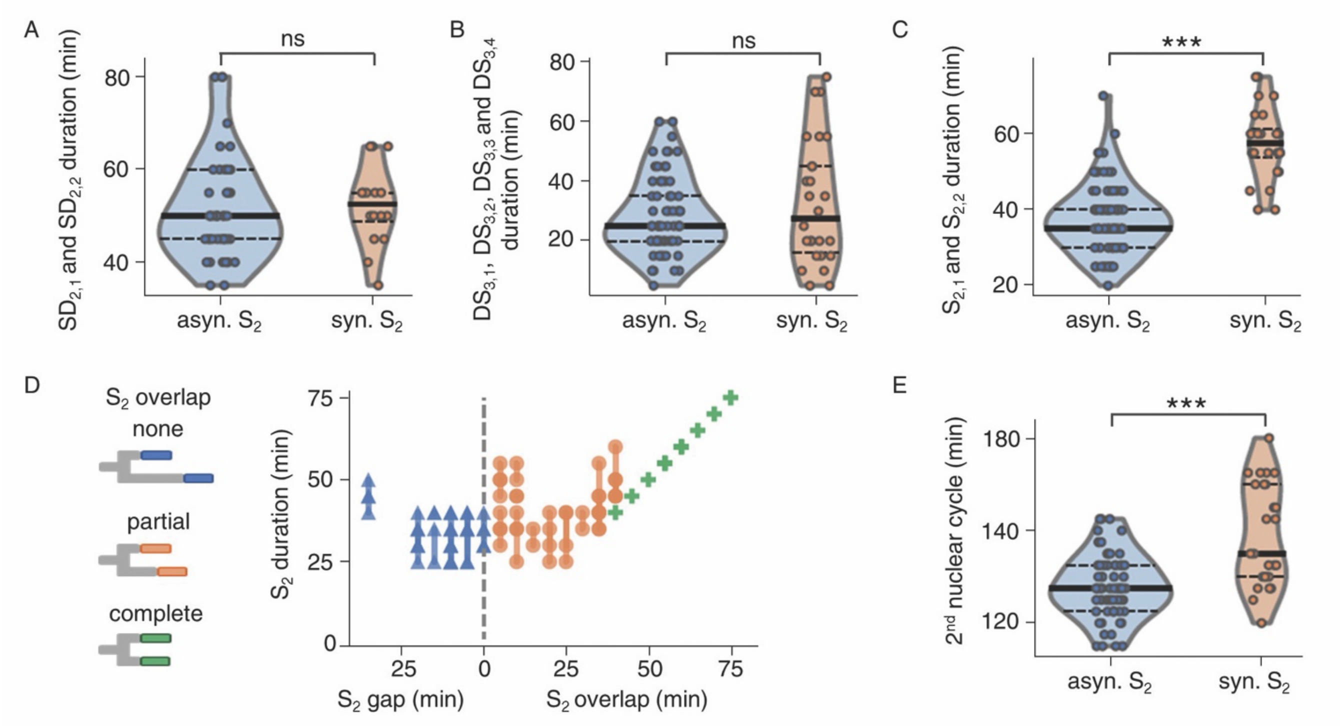


**Fig. S7. The effect of synchronous S-phases on nuclear multiplication dynamics in sister nuclei.** (A) The duration from S-phase to division (SD) was indistinguishable in sister nuclei with synchronous and asynchronous S-phases; solid lines, median; dashed lines, quartiles; two-sided Mann–Whitney *U* test, $p = 0.47$. (B) The duration from division to S-phase (DS) was indistinguishable in sister nuclei with synchronous and asynchronous S-phases; solid lines, median; dashed lines, quartiles; two-sided Mann–Whitney *U* test, $p = 0.68$. (C) Synchronous S-phases in pairs of sister nuclei were significantly longer; solid lines, median; dashed lines, quartiles; two-sided Mann–Whitney *U* test, $p = 3.4e^{-11}$. (D) The duration of S-phases in pairs of sister nuclei (vertical lines) *versus* the temporal gap or overlap between S-phases; asyn., asynchronous; syn., synchronous. (E) The second nuclear cycle was significantly longer in pairs of sister nuclei with synchronous S-phases; solid lines, median; dashed lines, quartiles; two-sided Mann–Whitney *U* test, $p = 2.36e^{-05}$.

**Table S1. Parameters of the Gamma distribution used in the branching process models, based on time-lapse microscopy of the initial nuclear cycles.** The shape and rate parameter of the Gamma distributions, α and β, were estimated by maximizing a log-likelihood function. The KS test is performed as a one-sample test with a two-sided alternative hypothesis.

| | **First nuclear cycle** | | **Second and higher nuclear cycle** | |
|---|---|---|---|---|
| **Nuclear cycle phase** | $S_1$ | $D_1$ | $S_{>1}$ | $D_{>1}$ |
| **Input data** | $S_1$ | $SD_1+DS_{2,1}$, $SD_1+DS_{2,2}$ | $S_{2,1}$, $S_{2,2}$ | $SD_{2,1} + DS_{3,1}$, $SD_{2,1} + DS_{3,2}$, $SD_{2,2} + DS_{3,3}$, $SD_{2,2} + DS_{3,4}$ |
| $\boldsymbol{\alpha}$ | 8.14 | 18.77 | 12.49 | 25.85 |
| $\boldsymbol{\beta}$ | 0.15 | 0.15 | 0.30 | 0.32 |
| **KS test *p*** | 0.24 | 0.033 | 0.013 | 0.17 |

**Movie S1.**
Sections of an electron tomogram of a high-pressure frozen multinucleated *P. falciparum* blood-stage parasite; resolution reduced by a factor of 4; every $4^{th}$ slice is shown; brightness and contrast adjusted for better visibility; bar, 1 $\mu$m.

**Movie S2.**
Sections of an electron tomogram of a high-pressure frozen multinucleated *P. falciparum* blood-stage parasite; resolution reduced by a factor of 4; every $8^{th}$ slice is shown; brightness and contrast adjusted for better visibility; bar, 1 $\mu$m.

**Movie S3.**
Time-lapse microscopy of a reporter parasite expressing nuclear mCherry (magenta) and stained with the DNA dye 5-SiR-Hoechst (multicolor); DIC image, single slice; fluorescent images, z-projection of 9 z-slices; brightness and contrast adjusted for better visibility; bar, 2 $\mu$m.

**Movie S4.**
Time-lapse microscopy of a reporter parasite expressing nuclear mCherry (magenta) and PCNA1::GFP (green); images, z-projection of 17 slices; brightness and contrast adjusted for better visibility; bar, 2 $\mu$m.

**Movie S5.**
Time-lapse microscopy of a reporter parasite expressing nuclear mCherry (magenta) and PCNA1::GFP (green); images, z-projection of 17 slices; brightness and contrast adjusted for better visibility; bar, 2 $\mu$m.

**Movie S6.**
Time-lapse microscopy of a reporter parasite expressing nuclear mCherry (magenta) and PCNA1::GFP (green) showing synchronous S-phases (i.e. nuclear accumulation of PCNA1::GFP); images, z-projection of 17 slices; brightness and contrast adjusted for better visibility; bar, 2 $\mu$m.
